# Effect of thermal and high-pressure processing on the thermo-rheological and functional properties of common bean (*Phaseolus vulgaris* L.) flours

**DOI:** 10.1101/2020.03.10.986117

**Authors:** Tiantian Lin, Cristina Fernández-Fraguas

**Author notes:** Corresponding Author: Dr. Cristina Fernández-Fraguas, Human & Agricultural Biosciences Bldg.1, Room 402B, 1230 Washington St. SW, Blacksburg, Virginia 24061, USA.

## Abstract

The effect of hydrothermal (HT) (boiling for 15 or 120min) and high-hydrostatic pressure (HHP) (150, 300, 450, and 600MPa for 5, 10 or 15min) processing on the rheological, pasting, thermal and functional properties of bean flours was investigated. HT and HHP treatments differently affected these properties. HT120 led to maximum values of elastic and viscous moduli (*G*′, *G*″), and gel strength of bean flours. HHP enhanced *G*′, *G*″ and gel strength as the pressure and holding time increased. The viscoelastic properties of HT120 and HHP600/5-treated bean flours correlated with the increased viscosity of these samples. The pasting profiles and thermograms indicated a full, partial, and limited starch gelatinization in HT120, HHP600/5 and HHP ≤ 450MPa samples, respectively. Enthalpy values showed that HT120 caused a higher degree of protein denaturation than HHP, with protein denaturation increasing as pressurization and time increased. This had an impact on protein solubility and emulsifying activity of flours which were significantly diminished by HT15/HT120, but maintained or slightly decreased by HHP. Nevertheless, HHP-treated samples showed enhanced emulsifying stability with increased pressure and holding time. These results demonstrate that HHP has the technological potential to manufacture bean flours with a range of functionalities into diverse food products.

## 1. Introduction

Dry or common beans (*Phaseolus vulgaris* L.), one of the most important legume crops worldwide, are valuable and inexpensive source of functional food ingredients. Due to their high content in fermentable and low-digestible carbohydrates as well as micronutrients (Ai, Cichy, Harte, Kelly, & Ng, 2016; Hall, Hillen, & Garden Robinson, 2017), consumption of dry beans offers an underexploited solution to fight against several cardiovascular disorders (Padhi & Ramdath, 2017). In addition, due to their high content in protein, common beans are promising ingredients for developing meat-substitutes and gluten-free food products of high nutritional value (Foschia, Horstmann, Arendt, & Zannini, 2017). However, despite food nutritional quality has gained a significant importance among consumers, consumption of dry beans per capita has seen a slow but steady decline in developed countries (Siddiq & Uebersax, 2012); specifically, 80% of the population do not consume the recommended daily intake of dry beans (USDA, 2015). There is, therefore, an opportunity for the food industry to exploit the health benefits of common beans in the development of attractive, new, ready-to-use bean ingredients. The use of common beans in the form of flours or pastes could increase their application range and stimulate bean consumption while improving the nutritional quality of a variety of processed food products.

The development of bean-based foods requires characterization of the physico-chemical and functional properties of bean flours as model systems before their incorporation to food products. In this regard, the rheological and thermal behavior of bean dispersions are of major importance when designing and modelling manufacturing processes of semi-solid foods. In addition, the viscosity and viscoelastic properties of bean dispersions can play a role on the textural and sensorial attributes of formulated foods (J. Ahmed, Ptaszek, & Basu, 2017), and therefore are of practical significance for acceptance and consumption. Both, the composition (i.e. proteins, starch, cell wall polysaccharides and minor components) and the microstructure of bean flours, contribute to these properties and functionalities (Kaur & Singh, 2005).

Processing technologies can diversify the use of bean flours as ingredient in manufactured food products by altering their functional properties. Hydrothermal (HT) processing, such as boiling and canning, is widely used to manufacture beans into edible foods. However, high cooking temperature and/or prolonged heating time has reportedly caused negative changes in the sensorial quality and nutritional value of legumes (Lee et al., 2018). The concern about ultra-processed food and health outcomes that has grown in recent years, has driven the need for development and validation of new generation processing technologies. High-hydrostatic pressure (HHP) is a non-thermal technology that has been mainly applied to increase the microbiological safety of foods during the last decade (Xu, Lin, Wang, & Liao, 2015). This technology has also shown encouraging potential to manipulate the physicochemical properties of plant foods with minimum loss of sensory and nutritional quality (Zou et al., 2016). Specifically, HHP has shown to modify the conformational arrangement of proteins affecting protein functionalities such as protein solubility, water/oil binding ability and emulsifying properties (M. D. Alvarez et al., 2014). In addition, gelatinization and gelation of starch, a primary component of beans, occurs during pressure treatment, and the extent of these phenomena depends on the starch type, pressure level, time and temperature of pressurization, water amount and matrix surrounding the starch. Hence, in contrast to most processing techniques, HHP can be easily applied to obtain a range of functionalities by controlling pressure conditions. Most of the studies found in the literature are focused on the effect of HHP on the properties of isolated starch and protein from pulses (Jasim Ahmed et al., 2018; Garcia-Mora, Peñas, Frias, Gomez, & Martinez-Villaluenga, 2015) or chickpea, lentil and pea flours (Jasim Ahmed, Varshney, & Ramaswamy, 2009; Leite, de Jesus, Schmiele, Tribst, & Cristianini, 2017). Additionally, most of these studies were performed by applying thermal or high-pressure treatment to the whole seeds before applying mechanical treatment. However, to our knowledge, a comprehensive physico-chemical characterization of HHP-processed flour systems from pinto beans, the most highly consumed type of bean in the US (Gittlein, 2018), is not described in the literature. Therefore, this study aimed to develop pinto bean flours using mechanical and HHP processing, and to systematically compare how HT and HHP treatments affect the rheological, thermal, pasting and resulting functional properties of bean flours.

## 2. Materials and Methods

### 2.1 Materials

Pinto beans were chosen as representative common variety of the market class of *Phaseolus vulgaris*. Raw bean seeds were donated from ADM (Archer Daniels Midland Company, IL, USA) and stored at room temperature. Beans were grown in different regions of North Dakota and harvested in the same crop year. All the lab chemicals were purchased from Fisher Scientific (Hampton, USA).

### 2.2 Hydrothermal and High-Hydrostatic Pressure treatments

The preparation of bean samples was done as described previously (Lin, O’Keefe, Duncan, & Fernández-Fraguas, 2019) with the incorporation of an additional Hydrothermal (HT) treatment. Briefly, whole beans were ground by a grinder (Waring commercial, USA) and the flour had a mean particle size in the range 450∼550 *µ*m as measured by a LS13320XR laser diffraction particle size analyzer (Beckman Coulter, Inc., USA). The effect of High-hydrostatic pressure (HHP) was studied on flours, as a function of pressure level (150, 300, 450 and 600 MPa, 25°C) and time (5, 10 and 15 min) at a 1:2 flour-to-water ratio. Treatment at 600MPa was only operated for 5min due the maximum capacity limit of the equipment. HT treatment consisted of a cooking treatment in boiling water at atmospheric pressure for 15min or 120min. The convention for naming samples is to list the type of processing method, with the remainder being the processing conditions (pressure and/or time). For example, HHP-treated samples at 150MPa for 5min is referred to as HHP150/5, and HT-treated samples for 15min is referred to as HT15.

### 2.3 Rheological characterization

Rheological properties of bean samples were determined using a DHR3 rheometer (TA instruments, Waters Co., UK) using a plate-plate measuring system (40mm diameter, 1mm gap) and a solvent trap to minimize moisture loss. The test temperature was set at 25 °C by using a circulating water system. An amount of sample was loaded into the geometry and allowed to rest for 5min. The viscoelastic behavior of samples was studied using small-amplitude oscillatory shear (SAOS) measurements. Amplitude sweeps at a constant angular frequency (*ω*) of 1 rad/s and shear strain increasing from 0.001 to 10 (0.1%-1000%) were initially performed to determine the linear viscoelastic region (LVR) of samples. Frequency sweeps were then performed from 0.1 to 100 rad/s at a constant shear strain within the LVR. The evolution of the elastic modulus (*G’*), viscous modulus (*G’’*) and complex viscosity (η^*^) with frequency was determined. A power law model (Eq. (1) and Eq. (2)) was used to fit frequency sweep data:

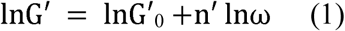

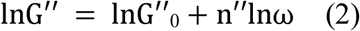

Where G^’^_0_ and G^’^_0_ are related to the strength of the network and, n’ and n” provide insight into the viscoelastic behavior. The value of (G′_0_ − G″_0_) was used to evaluate gel strength (Lapasin, 2012). Steady shear measurements were carried out at shear rate varying from 0.1 to 120 1/s. Data were fitted to different rheological models including Newtonian, Bingham, Power Law, and Herschel-Bulkley. The Herschel-Bulkley model (Eq. 3) showed the best fitting, and was therefore used to describe the flow behavior of bean dispersions:

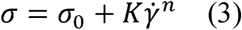

Where *σ* is shear stress (Pa), *σ*_0_ is yield tress (Pa), K is consistency coefficient (Pa s^n^), 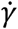 is share rate (1/s), and n is flow behavior index.

### 2.4 Pasting properties

Pasting properties were determined using the same rheometer, geometry and gap as described in section 2.3. A temperature (heating + cooling) ramp was performed from 30-95 °C, and from 95°C-30 °C, respectively, at a rate of 5°C/min, and at constant *ω* (1 rad/s) and stress (25 Pa). Silicon oil and solvent trap were used to prevent the evaporation of water. The pasting profile of samples was obtained by monitoring the evolution of the complex viscosity η* with temperature, and the peak viscosity, heat-paste viscosity and cold-paste viscosity of bean samples were recorded.

### 2.5 Thermal properties

A multicell differential scanning calorimeter (MC-DSC, TA Instruments, USA) was used to determine the thermal properties of samples. Samples were prepared by mixing bean flour with water (1:2 w/v) at room temperature. 150±5mg sample was weighed directly into Hastelloy ampoules and sealed. D/I water in the same range of weight was used as baseline of heat change in each ampoule. An empty ampoule was used as reference. The instrument was equilibrated for at least 60min before starting the heating/cooling scanning cycle from 25 °c-150 °C at 2°C/min. Thermal parameters including onset (T_o_), peak (T_p_) and conclusion (T_c_) temperature, as well as enthalpy change (*ΔH*) of the transition were calculated using the Nanoanalyze software (TA Instruments).

### 2.6 Functional properties

#### 2.6.1 Protein solubility

Protein solubility was measured by the Bradford method (Bradford, 1976), according to the method of (Boye et al., 2010) using the Coomassie Brilliant Blue G-250 reagent and a GENESYS™ 10S UV-Vis Spectrophotometer. Protein solubility was calculated as the ratio of soluble protein in the supernatant to that of total protein in the original mixture dispersions.

#### 2.6.2 Water-holding capacity and Oil-binding capacity

The water-holding capacity (WHC) and oil-binding capacity (OBC) of samples were determined according to (Setia et al., 2019). The WHC and OBC were calculated as the weight of retained water or bound oil (g), respectively, by sample (g).

#### 2.6.3 Emulsifying properties

The emulsifying properties of bean flours were analyzed according to the method of Pearce & Kinsella, 1978 and Shen, Fang, Gao, & Guo, 2017 by the turbidimetric technique, and were expressed as EAI (Emulsifying activity index,m^2^/g) and ESI (Emulsifying stability index, min). EAI measures the maximum surface area occupied by surface active molecules and their capacity to form an emulsion, while ESI measures the stability of an emulsion over a specific time (Kiosseoglou & Paraskevopoulou, 2011).

### 2.7 Data and statistical analysis

All the tests were performed in triplicate and values were reported as means and standard deviation. The data were analyzed using Analysis of Variance (ANOVA). The Tukey HSD comparison test were conducted to evaluate the significant differences among experimental mean values at the significance levels of 0.05 (*p*<0.05). All the statistical analyses were conducted and plotted by JMP Pro13 (SAS Institute, USA) and Graphpad Prism8 (GraphPad Software Inc., USA).

## 3. Results and discussion

### 3.1 Rheological characterization of bean flour dispersions

#### 3.1.1 Viscoelastic behavior

The frequency-dependent behavior of raw, HT- and HHP-treated samples is shown in Fig.1. Control and processed samples showed a predominantly more elastic response with magnitudes of *G’* higher than *G’’* through the angular frequency range studied. Both moduli were relatively independent of the oscillation frequency without crossover of *G*′ and *G*″, indicating that bean dispersions can be characterized as weak gels (Rao, 2010). Similar weak gel-like behavior was reported in thermal-treated chickpea flours (M. Alvarez, Fuentes, & Canet, 2015) and HHP-treated lentil flour slurries (Ahmed, Varshney, & Ramaswamy, 2009). Frequency sweep data were fitted by Eqs. (1) and (2) and the obtained magnitudes of estimated slopes (*n’* and *n’’*) and intercepts (*G’’*_*0*_ and *G’*_*0*_) also support the weak gel character observed (Table 1). While slopes provide insight into the viscoelastic and gelling behavior, the intercepts are related with the network strength. All bean dispersions had similar magnitudes of slope *n’* (0.11-0.14) and thus a similar frequency dependence of *G*′, implying that the weak gel character of bean dispersions was not modified by thermal or pressure processing. However, both processing treatments had a significant effect on the network strength, with oscillatory data indicating an increase of rigidity of the gels after processing.

After HT treatment, both HT15 and HT120-treated samples exhibited a significant increase in both *G*′ and *G*″ moduli (Fig. 1A) reflecting the creation of a more solid-like structure compared to untreated samples. HT15 and HT120-treated beans showed also the highest values of intercepts (*G*′_*0*_ and *G*″_*0*_) as well as gel strength (*G’*_*0*_ *-G’’*_*0*_) (Table 1), supporting the fact that a high entanglement density three-dimensional network of cross-linked protein and starch is formed when gelatinization occurs during thermal treatment. It is also possible that an increased stiffness of the swollen starch granules consequence of starch retrogradation occurred during storage, contributing to the large gel strength observed (Keetels, Vliet, & Luyten, 1995). All HHP treatments increased *G*′ and *G”*, with both moduli gradually increasing with increasing pressure (Figs. 1A & B) and holding time (Fig. 1C). This might indicate an increase in molecular interactions and a strengthening of bean microstructure compared to raw samples, which was in agreement with previous findings reported in flour dispersions (Ahmed et al., 2009). Values of intercepts and gel strength revealed differences between pressure levels. Whereas low pressures (150 MPa) were not enough to cause any significant change in the network strength of bean dispersions at any of the holding times, higher pressures (600 MPa) resulted in the formation of a stronger network dominated by the presence of a cross-linked arrangement of partially gelatinized starch and/or denatured protein, even at short holding time (Jiang, Li, Hu, Wu, & Shen, 2015). Pressure-induced starch gelatinization seems to occur over a range of pressures and a critical level of pressure needs to be reached for gelatinization to occur effectively (Bauer & Knorr, 2005; Oh et al., 2008). During treatment at low pressures, the compression force exerted on starch granules might predominate over granule swelling; as the pressure increases the granule rigidity increases but also more water penetrates into the granule promoting swelling and gelatinization (Jiang et al., 2015). Particularly, HHP 600MPa/5 min resulted in the formation of network structures of slightly lower strength to that created after HT treatment. The effect of HHP on complex viscosity (η^*^) (Fig. 1A) clearly shows an increase in viscoelasticity with increased pressure. η^*^ values of HT120- and HHP600/5-treated samples were very similar and significantly higher than the corresponding to raw samples which supports the more viscoelastic character of processed gels under these conditions. Overall, the greater viscoelastic moduli, η^*^ and gel strength observed in HT- and HHP600/5-treated beans indicated molecular interactions and major structural modifications under those conditions.

**Table 1.**
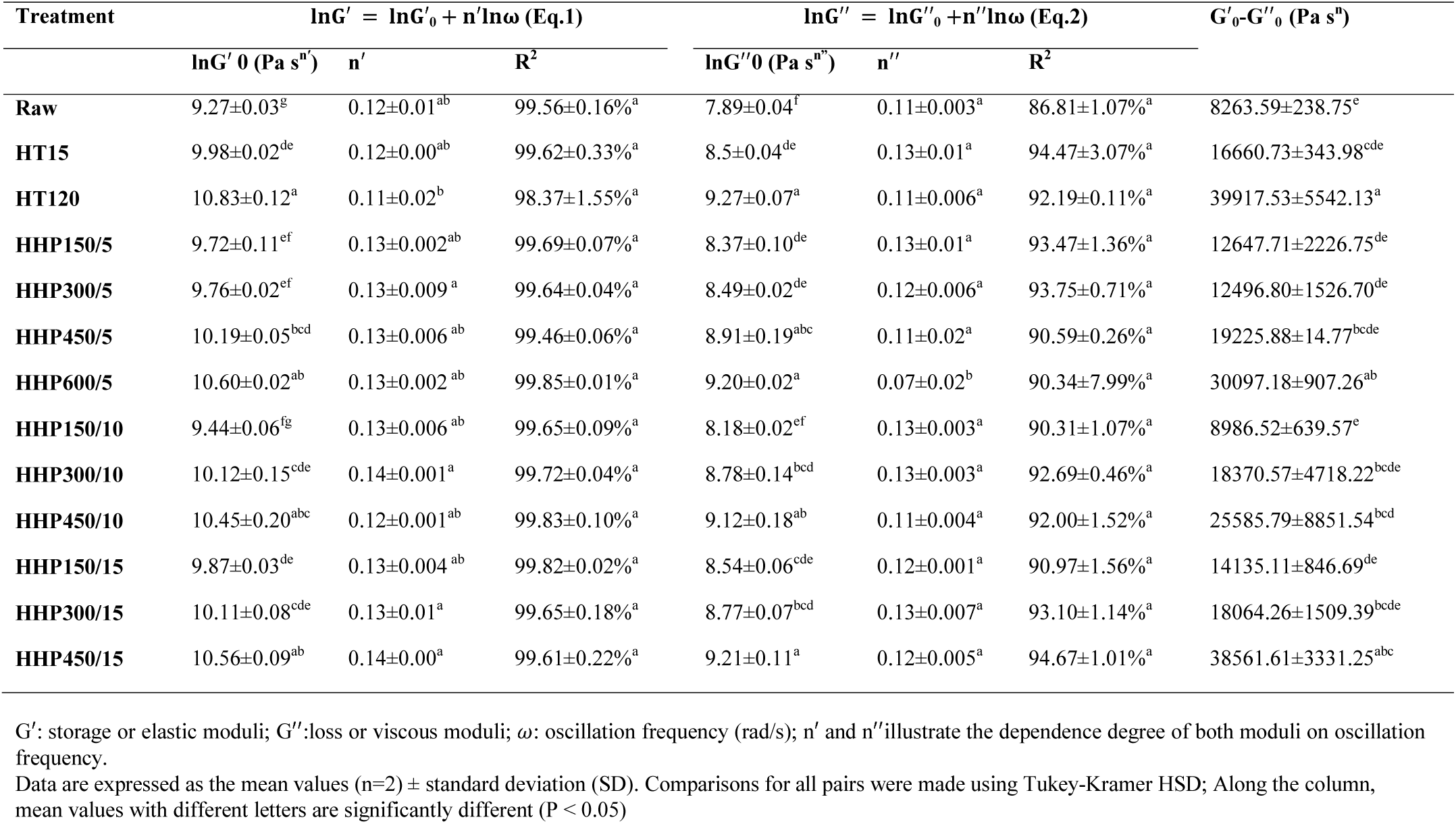
Power law parameters of Eq. (1) and (2) from mechanical spectra of bean flour dispersions treated by hydrothermal and high-hydrostatic pressure processing.

**Fig. 1.**
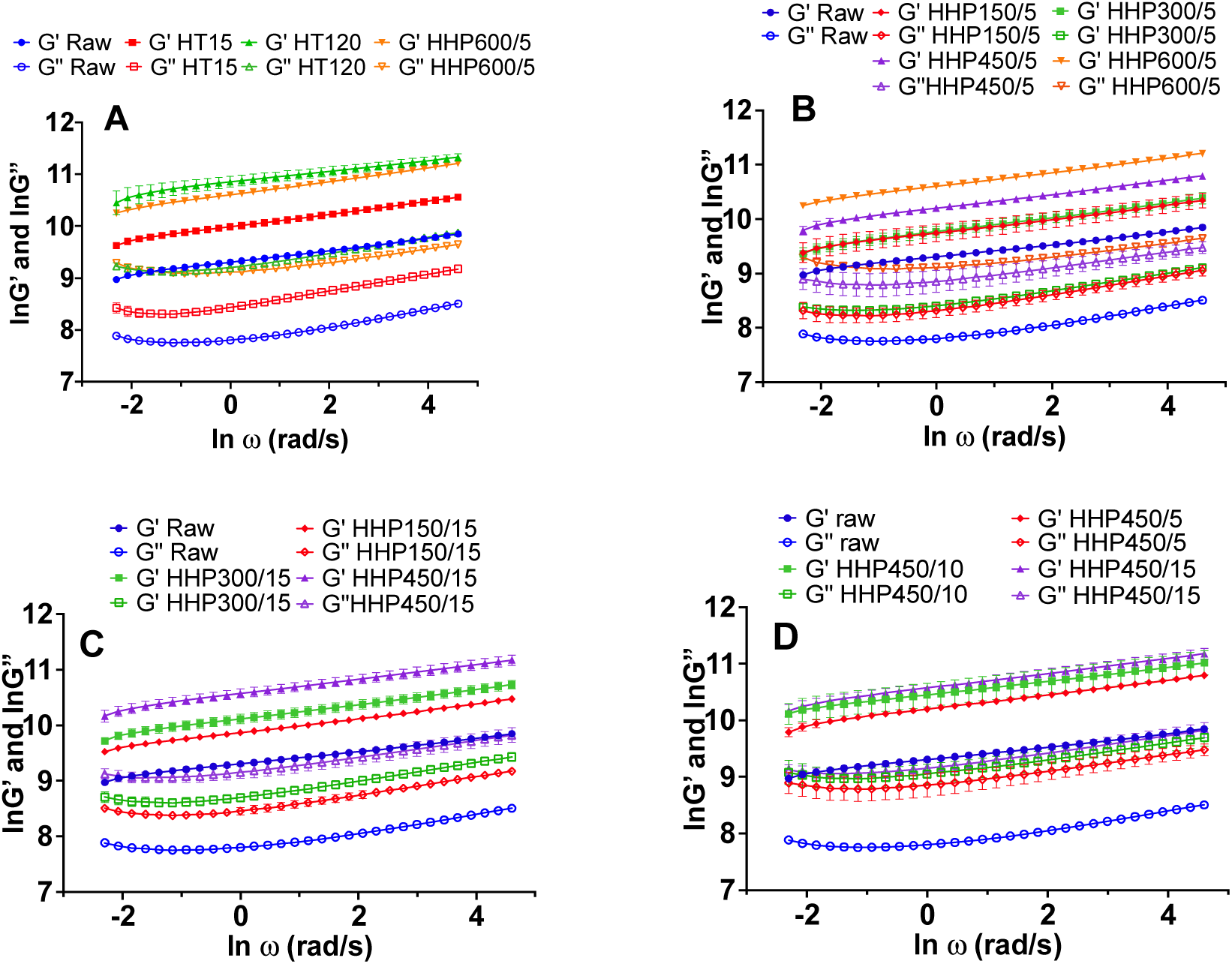
Viscoelastic properties of bean flours treated by hydrothermal and high-hydrostatic pressure processing as a function of (A) treatment type, (B) treatment pressure at 5min, (C) treatment pressure at 15min and (D) treatment time at 450 MPa. G’: storage or elastic moduli; G”: loss or viscous moduli; *η*\*: complex viscosity Data are expressed as the mean values (n=3) ± standard error (SE) bar

#### 3.1.2 Flow behavior

Fig. 2 shows the flow behavior of raw, HT- and HHP-treated bean samples. The apparent viscosity (*η*_*a*_) of all bean samples decreased with increasing shear rate (*γ*), indicating a shear-thinning behavior, characteristic of most non-Newtonian foods (Rao, 2010). Similar shear-thinning behavior was also reported in previous studies on pulse dispersions (M. D. Alvarez, Herranz, Campos, & Canet, 2017). Among the flow models used in this study, the Herschel– Bulkley model (Eq. (3), which is widely used to describe pseudoplastic behavior, showed the best fitting of flow parameters in bean samples (R >99%) (Table S1). Therefore, only the fitting parameters (*σ, σ*_0_, K and n) obtained with this model are discussed (Table 2). All samples showed *n* values <1 confirming the shear-thinning behavior of bean dispersions. Raw and processed samples did not show a large variation of *n* (0.40-0.53), which were comparable to values reported in chickpea slurries (Alvarez et al., 2017). HT treatment had a significant effect on the flow behavior of bean dispersions (Fig. 2A). Compared to HHP, HT-treated samples were less stable to shear, representative of a greater orientation and disentanglement of polymer chains under shear. In particular, K exhibited a maximum value in HT120 samples, ∼25 times higher than for raw samples, in accordance with the highest *η*_*a*_ observed over the whole range of *γ*. This could indicate a significant extent of swollen starch granules, consequence of starch gelatinization. Regarding HHP treatments, the effect of pressure level was dependent on the length of treatment. Whereas at 5 min, *η*_*a*_ and *K* increased with increasing pressure (Fig. 2B), at 10/15 min, *K* decreased with increasing pressure (Fig. 2C), even if no clear effects on *η*_*a*_ were noticed. For a specific pressure (e.g. 450 MPa), *K* and *η*_*a*_ decreased with increased holding time (Fig. 2D). The decreased *K* and *η*_*a*_ could be due to the presence of pores on the surface that increased in size with increasing holding time, as we previously reported (Lin et al., 2019). Other studies also reported an increase of *K* with increasing pressure in mung starch (Jiang et al., 2015) and a decrease of *K* with holding time in chickpea dispersions (Alvarez et al., 2017). In any case, HT600/5 samples showed the highest *K* and *η*_*a*_ values within HHP-treated samples. Yield stress (*σ*_0_) is defined as the minimum shear stress required to initiate flow (Steffe, 1996). Raw samples showed a low *σ*_0_ that sharply and significantly increased after HT-treatment (Fig. 2A and table 2) reflecting the elevated resistance of heated bean dispersions to flow, which is consequence of the high cross-linked network that must be broken down (Duran & Costell, 1982; Qiu & Rao, 1988). Although *σ*_0_ values exhibited a non-identifiable trend with HHP treatment, 600MPa considerably decreased the *σ*_0_, which was consistent with studies on cereal starch (Li & Zhu, 2018). From these observations, it can be concluded that the rheological response of bean dispersions can be attributed to the degree of gelatinized starch and their cross-linking with proteins.

**Table 2.**
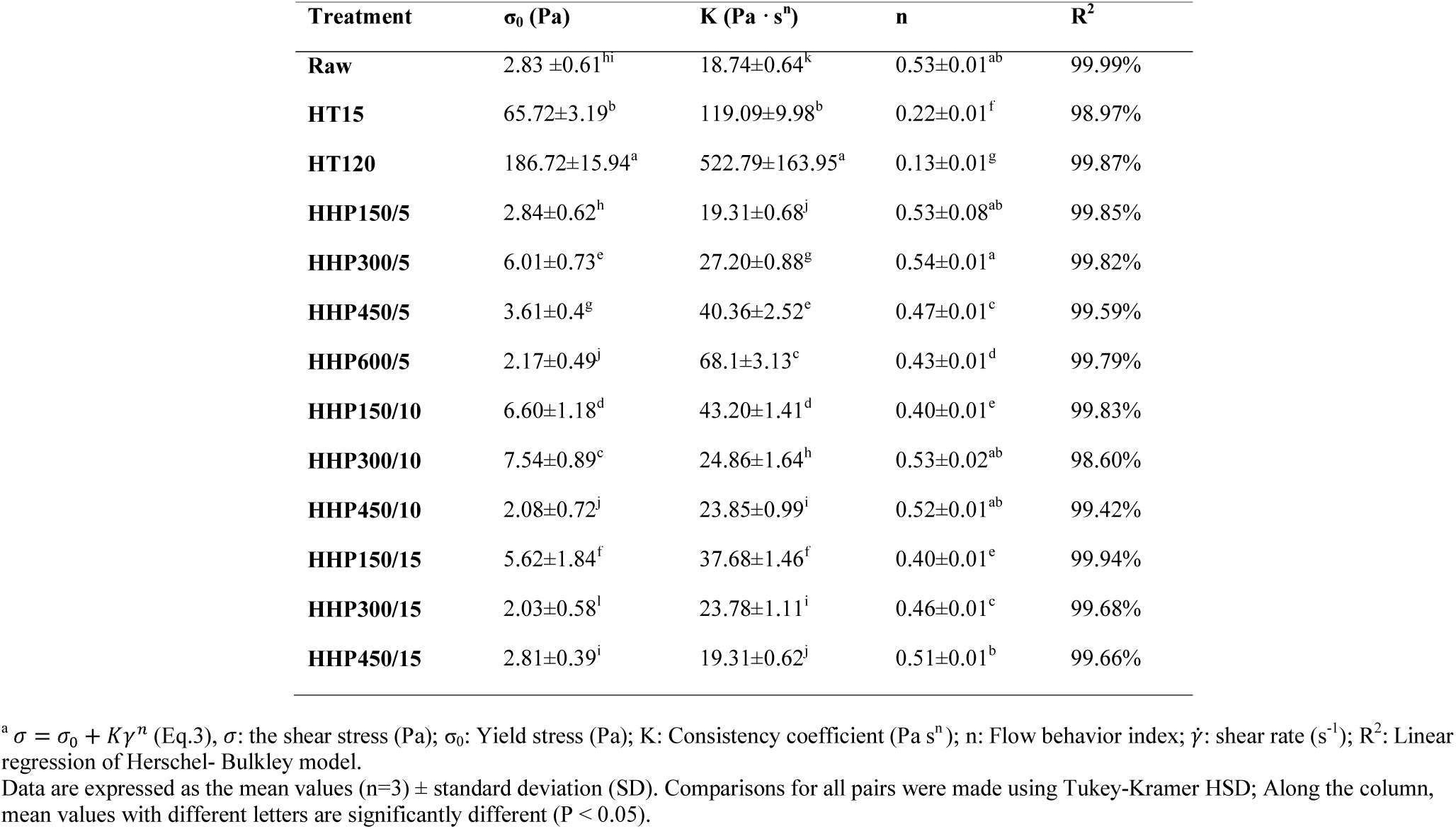
Flow behavior parameters of bean flour dispersions treated by hydrothermal and high-hydrostatic pressure processing estimated by the Herschel-Bulkley model ^a^

| **Treatment** | **$\sigma_0$ (Pa)** | **$K$ (Pa · s<sup>n</sup>)** | **n** | **R<sup>2</sup>** |
| --- | --- | --- | --- | --- |
| **Raw** | 2.83 ±0.61<sup>hi</sup> | 18.74±0.64<sup>k</sup> | 0.53±0.01<sup>ab</sup> | 99.99% |
| **HT15** | 65.72±3.19<sup>b</sup> | 119.09±9.98<sup>b</sup> | 0.22±0.01<sup>f</sup> | 98.97% |
| **HT120** | 186.72±15.94<sup>a</sup> | 522.79±163.95<sup>a</sup> | 0.13±0.01<sup>g</sup> | 99.87% |
| **HHP150/5** | 2.84±0.62<sup>h</sup> | 19.31±0.68<sup>j</sup> | 0.53±0.08<sup>ab</sup> | 99.85% |
| **HHP300/5** | 6.01±0.73<sup>e</sup> | 27.20±0.88<sup>g</sup> | 0.54±0.01<sup>a</sup> | 99.82% |
| **HHP450/5** | 3.61±0.4<sup>g</sup> | 40.36±2.52<sup>e</sup> | 0.47±0.01<sup>c</sup> | 99.59% |
| **HHP600/5** | 2.17±0.49<sup>j</sup> | 68.1±3.13<sup>c</sup> | 0.43±0.01<sup>d</sup> | 99.79% |
| **HHP150/10** | 6.60±1.18<sup>d</sup> | 43.20±1.41<sup>d</sup> | 0.40±0.01<sup>e</sup> | 99.83% |
| **HHP300/10** | 7.54±0.89<sup>c</sup> | 24.86±1.64<sup>h</sup> | 0.53±0.02<sup>ab</sup> | 98.60% |
| **HHP450/10** | 2.08±0.72<sup>j</sup> | 23.85±0.99<sup>i</sup> | 0.52±0.01<sup>ab</sup> | 99.42% |
| **HHP150/15** | 5.62±1.84<sup>f</sup> | 37.68±1.46<sup>f</sup> | 0.40±0.01<sup>e</sup> | 99.94% |
| **HHP300/15** | 2.03±0.58<sup>l</sup> | 23.78±1.11<sup>i</sup> | 0.46±0.01<sup>c</sup> | 99.68% |
| **HHP450/15** | 2.81±0.39<sup>i</sup> | 19.31±0.62<sup>j</sup> | 0.51±0.01<sup>b</sup> | 99.66% |

<sup>a</sup> $\sigma = \sigma_0 + K\dot{\gamma}^n$ (Eq.3), $\sigma$ : the shear stress (Pa); $\sigma_0$ : Yield stress (Pa); $K$ : Consistency coefficient (Pa s<sup>n</sup>); $n$ : Flow behavior index; $\dot{\gamma}$ : shear rate (s<sup>-1</sup>); R<sup>2</sup>: Linear regression of Herschel- Bulkley model.
Data are expressed as the mean values (n=3) ± standard deviation (SD). Comparisons for all pairs were made using Tukey-Kramer HSD; Along the column, mean values with different letters are significantly different (P < 0.05).

**Fig. 2.**
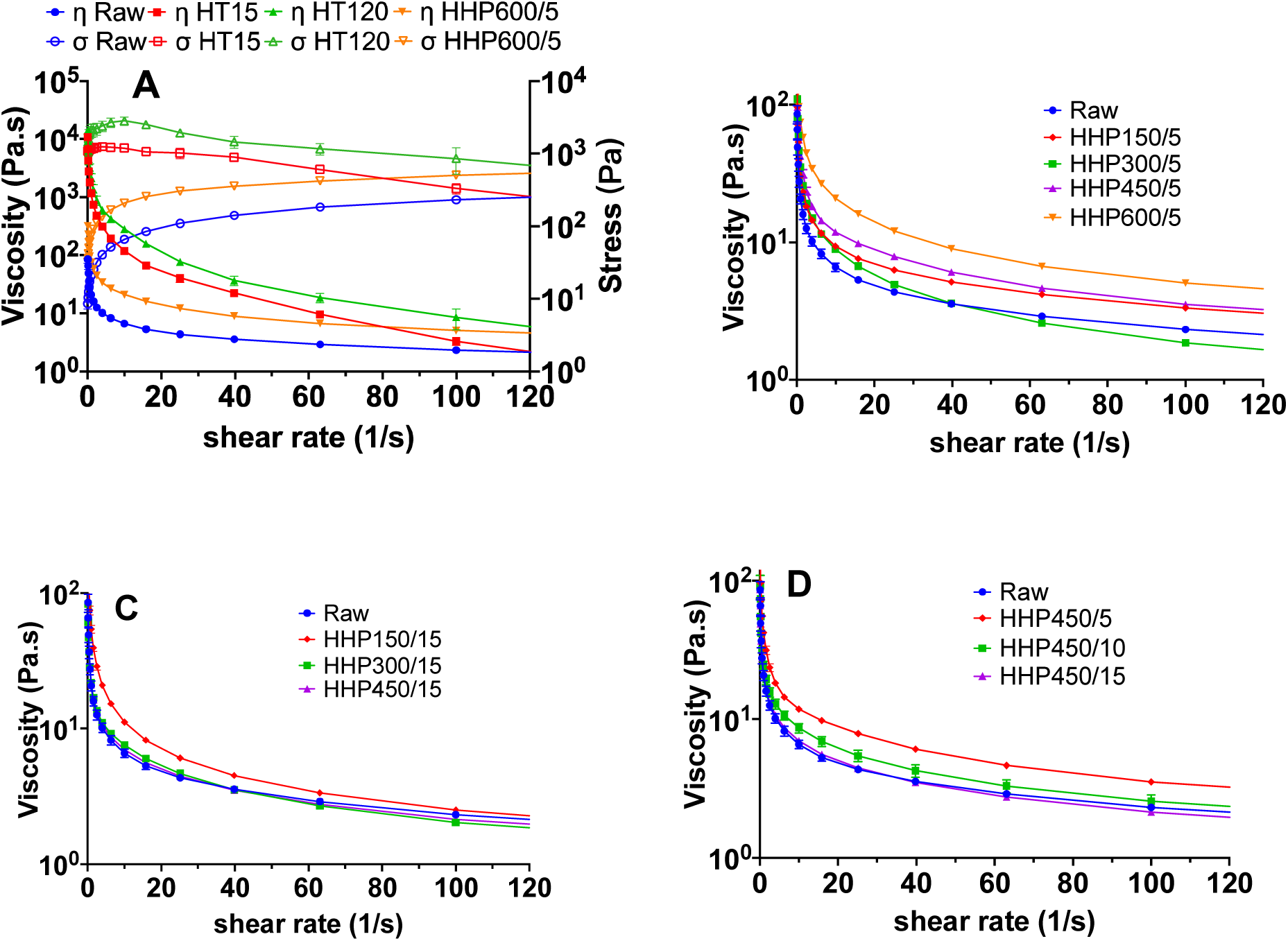
Flow behavior of bean flours treated by hydrothermal and high-hydrostatic pressure processing as a function of (A) treatment type, (B) treatment pressure at 5min, (C) treatment pressure at 15min and (D) treatment time at 450 MPa. Data are expressed as the mean values (n=3) ± standard error (SE) bar.

### 3.2 Pasting properties of bean flour dispersions

The pasting profile of bean dispersions is showed in Fig. 3, and the pasting parameters (i.e. initial viscosity, peak viscosity (PV), hot-paste viscosity (HV) and cold-paste viscosity (CV) are summarized in Table S2. Significant qualitative and quantitative differences in pasting properties were observed among raw, HT- and HHP-treated flour dispersions during heating and cooling. Raw samples initially showed a viscosity of ∼533 Pa.s that increased as the dispersion was heated and starch granules swelled. Starch granules were fully swollen and gelatinized at ∼74 °C reaching a maximum viscosity of 9970 Pa.s (i.e. PV). This gelatinization temperature is in the range of gelatinization temperature of pinto bean starch reported previously (Du, Jiang, Yu, & Jane, 2014; Rui & Boye, 2012). Pulse starch contains ∼30% amylose (∼10% higher than cereals starches) which is partly responsible for the high gelatinization temperature showed by common beans compared to cereals (∼65°C) (Rebello, Greenway, & Finley, 2014; Sharma & Gujral, 2010). As the temperature was further increased, a sharp decrease in viscosity was observed as consequence of the breakdown and disintegration of swollen granules, reaching hot paste values of ∼1869Pa.s at 85°C in raw samples (Fig.3A&TableS2). This drop in viscosity (∼70%) was more evident than the one observed in other pulses, such as pea and chickpeas (Ahmed, Thomas, Taher, & Joseph, 2016; Leite, de Jesus, Schmiele, Tribst, & Cristianini, 2017). Differences in the extent of viscosity breakdown among pulses could be due to differences in amylose content and extent of amylose leaching. Phosphorus content in starch has been also reported to contribute to the viscosity breakdown in different cultivars of pinto bean starch, as the repulsion between anionic phosphate groups could weaken amylopectin clusters, thus leading to a greater susceptibility of the starch granule towards shear (Ambigaipalan et al., 2011). Regarding HT treatment, due to the fact that the starch was pre-cooked, HT15 and HT120 samples showed a considerably high initial viscosity compared to raw (and HHP-treated) samples, which agrees with their flow behavior (Fig. 2A). In particular, it can be suggested that HT120 treatment led to a complete starch gelatinization since no PV appeared during heating cycle; however, HT15-treated samples showed a very small peak at ∼86-88°c which could suggest that a fraction of starch was not gelatinized (Figs. 3A&B). Regarding HHP, the higher initial viscosity and well-defined PV observed in HHP600/5-samples compared to raw samples indicates that these HHP conditions led to partial starch gelatinization. The rest of HHP samples showed a more similar profile, in terms of PV, to the raw samples (Figs. 3A, B, C &D) which suggested the presence of a low proportion (if any) of pre-gelatinized starch. However, both PV and HV were dependent on the pressurization level and time. While 600MPa increased PV, pressures≤450MPa increased gelatinization temperature and decreased PV values compared to raw samples. Specifically, PV decreased with decreasing pressure from 450-150MPa at holding time of 5min, but increased with decreasing pressure at the longer times (10 and 15min) (although the change was not significative between 300and 150MPa). These PV values could be partially explained by the presence of an increased number of small pores in the surface of bean dispersions pressurized at 150-450MPa (Lin et al., 2019) that contributed to the lesser capacity of starch granules to swell. On the other hand, the HV linearly increased with increasing pressure, which led to a reduced breakdown viscosity in all HHP samples (∼38-50%) compared to raw samples (∼70%) indicating that HHP enhanced the resistance to shear-thinning. This increased resistance of granules to rupture in HHP-treated samples may be related to a reinforced crystalline structure (Tester & Debon, 2000) and to the formation of a cross-linked network of starch-protein/fiber as consequence of high-pressure (Lin et al., 2019). The higher PV and HV showed by HHP600/5 compared to raw and HHP≤450MPa-treated samples (Fig.3B &Table S2), indicates that high-pressures applied for a short time can improve the thickening properties of bean flours or starch (Leite et al., 2017).

**Fig. 3.**
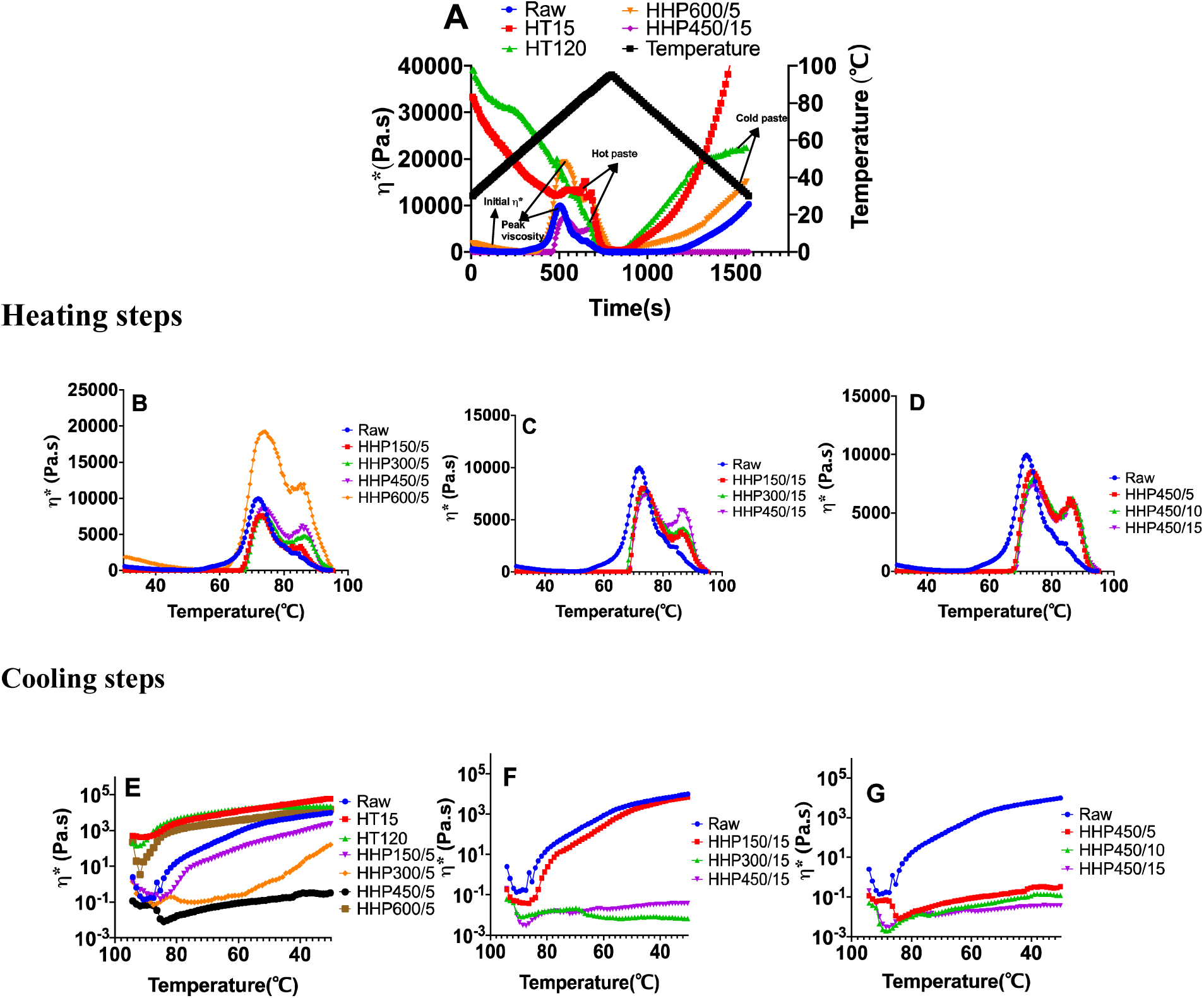
Effect of hydrothermal (HT) and high-hydrostatic pressure (HHP) treatments on the pasting profile of bean flour dispersions (A). Pasting properties of dispersions treated by HHP during 5min (B), 15min (C) and at 450 MPa (D) obtained during the heating cycle. Pasting properties of dispersions treated by HT (15 and 120 min) and by HHP during 5min (E), 15min (F) and at 450 MPa (G) obtained during the cooling cycle. Data are expressed as the mean values (n=3) ± standard error (SE) bar.

HT and HHP treatments also affected differently the pasting profile of dispersions during cooling (Figs.3A, E, F&G). The CV of raw sample was close to its PV values (Fig. 3A&Table S2). HT treatments increased the CV of bean dispersions (Fig. 3A&E), with the short-time treatment having a more pronounced effect on CV than the long one, indicating an increased starch retrogradation in HT15 samples. This different gelling capacity could be due to variations in the degree of granule breakdown, extent of gelatinization and hence HV, which was higher in HT15 samples. Regarding HHP, 600MPa increased CV values, whereas at HHP<600MPa/5min, CV dropped linearly with increasing pressure from 150 to 450MPa (Figs.3E&F). The decrease in CV was highly pronounced at 300-450MPa applied during longer times (Figs.3F&G). The presence of a higher proportion of gelatinized starch in HT and HHP600/5 samples compared to raw and the rest of HHP samples, consequence of a lower degree of starch crystallinity and packing, partly contributed to a greater extent of starch retrogradation, and consequently to a higher CV viscosity. The sharp reduction in CV observed at 300 and 450MPa (Figs.3A,E,F&G), could be attributed to the reinforced crystalline structure of starch in these samples and consequent lower amount of leached amylose. These lower pressures may also restrict the rearrangement/re-association of amylose molecules, which are the main contributor to starch retrogradation in short-term (Colussi et al., 2018). Similar effects of HT and HHP processing on the pasting profile of other pulses, such as mung beans and peas, were observed in previous studies (Kaur, Sandhu, Ahlawat, & Sharma, 2015; Leite, de Jesus, Schmiele, Tribst, & Cristianini, 2017).

### 3.3 Thermal properties of bean flour dispersions

The effect of HT and HHP processing on the thermal properties of bean flours is shown in Fig. 4. The T_0_, T_p_, T_c_ and *ΔH* corresponding to each endothermic peak are summarized in Table 3. Two endothermic thermal transitions are observed in the thermograms of all samples: Peak1 is associated to starch gelatinization and peak2, observed at higher temperature, is mainly related to protein denaturation and dissociation of lipid-starch complexes (Ahmed, Mulla, Arfat, & Kumar, 2017; Wright & Boulter, 1980). The observed T_p_ values in raw samples are similar to the T_p_ corresponding to starch gelatinization (∼84°C) and protein denaturation (∼98°C) reported for pinto bean (Chung, Liu, Peter Pauls, Fan, & Yada, 2008) and black/navy bean flours (Ai, Cichy, Harte, Kelly, & Ng, 2016). The beginning of protein denaturation (peak2 T_0_) coincided with the end of starch gelatinization (peak1 T_c_) as shown by the overlay of peak1 and peak2 in all bean samples (Fig.4). Both HT treatments significantly decreased the enthalpy values (*ΔH*) of raw samples (Fig.4A). At HT120, the bean starch completely gelatinized, and therefore, no glass transition temperature associated with gelatinization (T_p1_) was almost detected. Additionally, while HT120 significantly shift T_p1_ to lower temperatures, HT15 did not shift T_p1_, compared to raw samples. These results agree with the pasting (Figs.3A&B) and rheological properties (Fig. 2A) of HT15 and HT120-treated samples which confirm the partial and full starch gelatinization observed in thermal-treated samples during short and long time, respectively. Both, moderate and severe thermal treatment, did not significantly shift T_o,_ T_p_, T_c_ of peak2, however severe heating significantly decreased *ΔH* values (0.07J/g) indicating a high degree of protein denaturation and microstructural modification (Table 3).

**Table 3.**
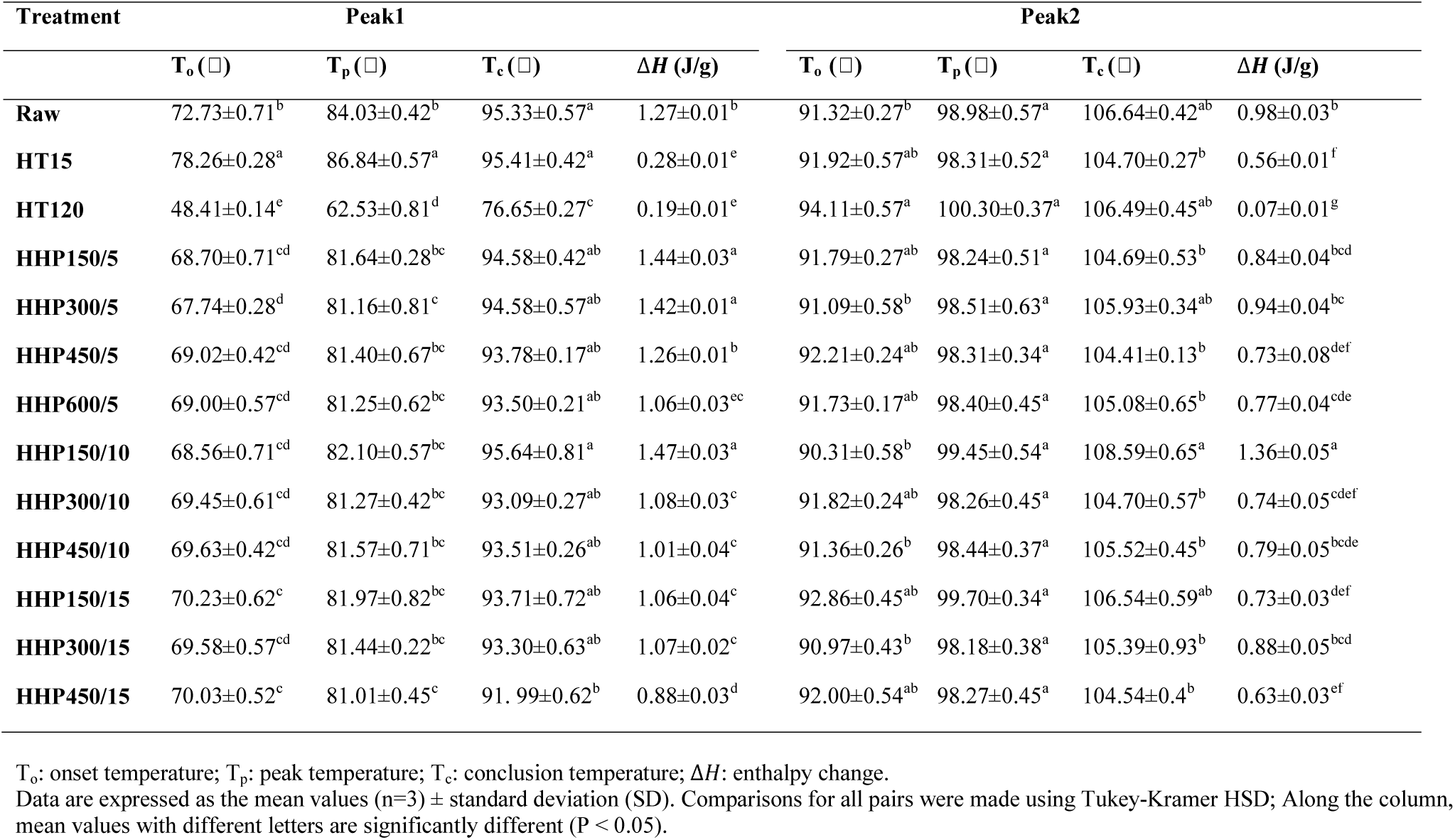
Thermal properties of bean flour dispersions treated by hydrothermal and high-hydrostatic pressure processing.

**Fig. 4.**
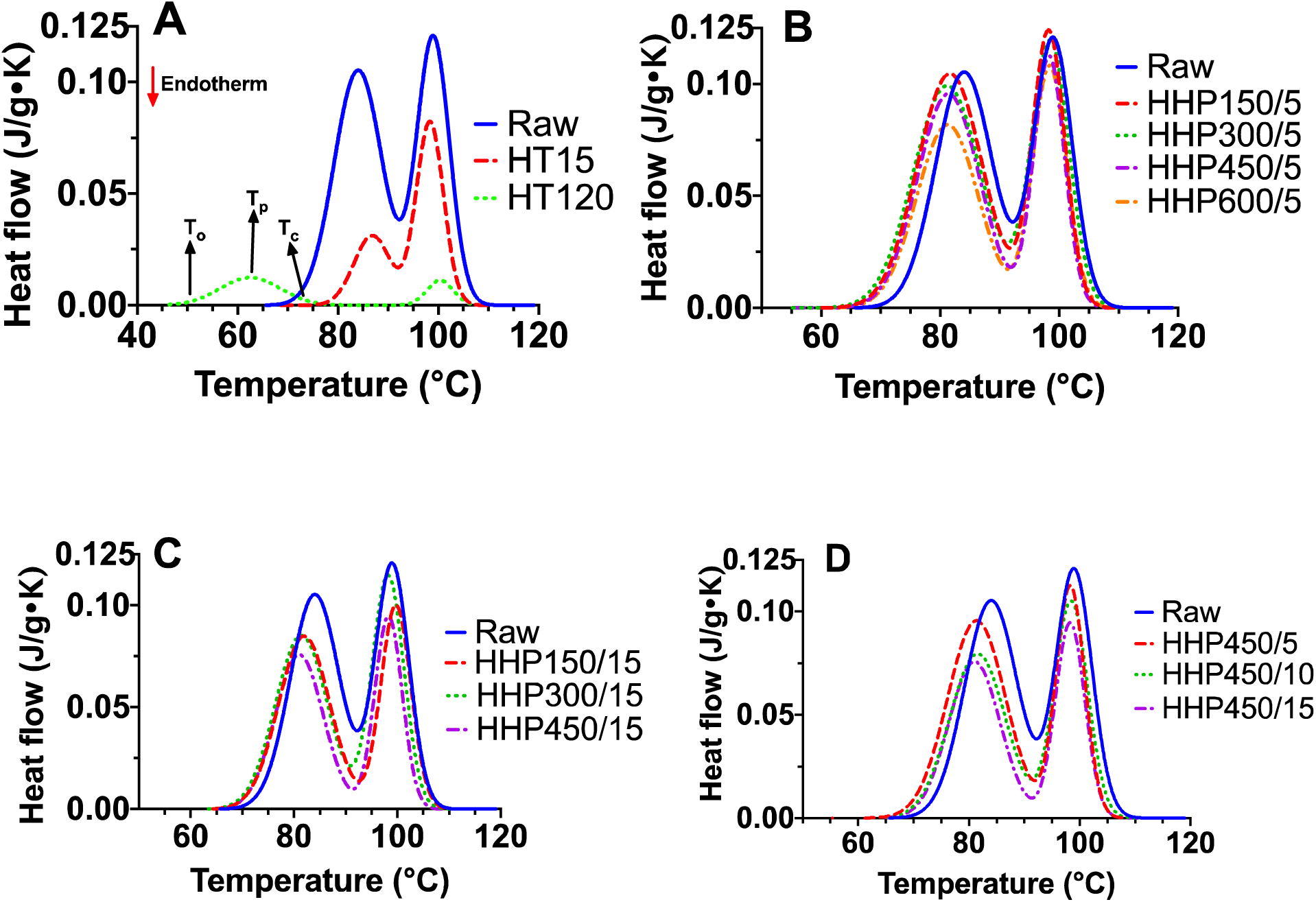
Thermal properties of bean flours treated by HT processing (A), by HHP processing for 5min (B), HHP processing for 15min (C) and HHP processing at 450MPa (D). Data are expressed as the mean values (n=3) ± standard error (SE) bar. T_o_: onset temperature; T_p_: peak temperature; T_c_: conclusion temperature.

Regarding HHP-treated samples, while T_p1_ was no significative different from raw samples (Figs.3A-D&Table 3), the enthalpy values gradually decreased as pressurization (Figs. 4B&C) and holding time (Fig. 4D) increased (Table 3), indicating an increased degree of starch gelatinization with pressure intensity. Similar trend has been reported by many researchers for starches (M. D. Alvarez, Fuentes, Olivares, & Canet, 2014; Leite et al., 2017)). As observed in HT samples, HHP had little influence on the transition temperature associated to peak2, whereas *ΔH* values dropped with increased pressure (Fig.4B&C) and holding time (Fig.4D), which could be related to both, breakup of non-polar interactions and aggregation upon unfolding of the native protein (Arntfield & Murray, 1981) as consequence of the elevated pressure and temperature. The extent to which aggregation affects enthalpy depends on the relative proportion of polar to non-polar amino acid residues exposed upon unfolding. Similar results were reported in isolated proteins from kidney beans pressurized at 200-600MPa 15min (Ahmed, Al-Ruwaih, Mulla, & Rahman, 2018). For both peak1 and peak2, HHP-treated samples showed significant higher *ΔH* than HT-treated samples, indicating that HHP leads to a lower extent of starch gelatinization and protein denaturation in bean flours (Table 3). The gelatinization of starch induced by HHP depends on the pressure level, holding time and the presence of an adequate amount of water (M.D. Alvarez et al., 2014) reported that complete starch gelatinization of chickpea slurries only occurred at 600MPa applied15min at low concentrations of flour (1:5 flour:water ratio). The HHP conditions used in the current study (i.e. 600MPa but shorter holding time and high concentrations of flour) may not be enough to cause complete gelatinization of bean starch, which is in accordance with previous studies on legumes using similar HHP conditions (Ahmed et al., 2016, 2009).

### 3.4 Functional properties of bean flours

#### 3.4.1 Protein solubility

The protein content in raw bean flour was 25.35%, and no significant differences were found with respect to processed samples (data not shown). Bean proteins are mainly storage proteins with globulins, considered as “multi-subunits” molecules of high molecular weight and relatively hydrophobic, constituting the main fraction (Rebello et al., 2014). The albumin fraction is of low-medium molecular weight and have a hydrophilic surface that renders this protein fraction water-soluble (Kiosseoglou & Paraskevopoulou, 2011). The effect of HT and HHP treatments on bean protein solubility is shown in Fig. 5. The protein solubility of raw pinto bean flours was ∼61%, similar to other pulse flours (Boye et al., 2010; Setia et al., 2019). HT treatment applied for 15min or 120min dramatically decreased the protein solubility by ∼80%. Similar results were reported in boiled chickpea and lentil flours 1h (Ma et al., 2011). The decreased protein solubility of HT-treated beans supports the observed thermal properties of these samples and could be a consequence of protein denaturation during heating which alters the hydrophilicity and hydrophobicity of the globulins and albumins surface in contact with the surrounding water. It is likely that a larger number of surface hydrophobic patches led to hydrophobic interactions over electrostatic repulsions between charged proteins, which induced protein aggregation and eventually lead to protein precipitation. It is also possible that cross-links between protein and starch molecules induced by heating led to the formation of aggregates, thus decreasing protein solubility (Ma et al., 2011). On the other hand, HHP treatments affected protein solubility in a different manner. Low pressures applied for short times (i.e. 5/10min) did not apparently affect protein solubility, whereas pressures >150MPa even if applied 5min significantly decreased protein solubility. Protein solubility was also reduced when low pressurization levels were applied for longer times. Similarly, Chapleau & de Lamballerie-Anton, 2003 reported that the protein solubility of lupin was only decreased at pressures>400MPa. These results agree with the higher degree of protein denaturation observed at either high pressure (600MPa) or long holding times (15min) (Figs.4B&D,Table 3).

**Fig. 5.**
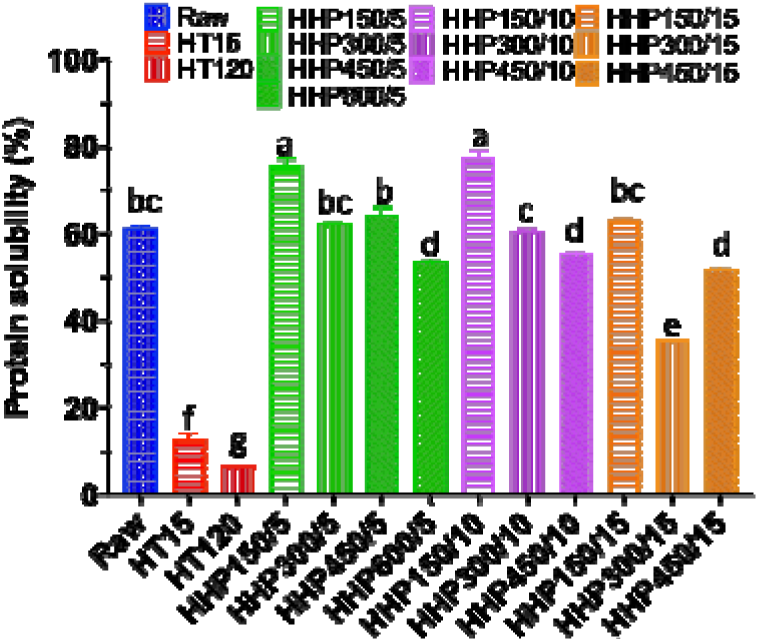
Protein solubility of bean flours treated by hydrothermal and high hydrostatic pressure processing. Data are expressed as the mean values (n=3) ± standard error (SE) bar.

#### 3.4.2 Water holding capacity and oil binding capacity

Fig. 6 shows the WHC and OBC of raw, HT- and HHP-treated bean flours. The WHC of raw bean flours was ∼1.7g/g (Fig. 6A) which was in accordance with previous studies (Deshpande, Sathe, Cornforth, & Salunkhe, 1982). The effect of HT-treatment on the WHC of bean flours was dependent on the duration of heating. HT120 significantly increased WHC, whereas HT15 showed no differences compared to raw flours. Similarly, other studies reported a significant increase of WHC of pulse flours after boiling 1h (Aguilera, Esteban, Benítez, Mollá, & Martín-Cabrejas, 2009). The increased WHC in HT120 samples could be partially related to protein denaturation while the less degree of protein denaturation observed in HT15 samples (Fig.4A) may be not sufficient to show effect on their WHC (Peyrano, Speroni, & Avanza, 2016). Heat-induced starch gelatinization could also increase water-absorption due to the formation of more disordered crystalline structures at extended heating times (Aguilera et al., 2009). Regarding HHP treatments, 5min had no significant effect on the WHC of bean flours, even at 600MPa. However, longer holding times at 300/450MPa significantly decreased WHC which could be explained by the decreased viscosity observed in these samples (Figs. 2C&D) (Ai et al., 2016). The presence of enlarged pores on the surface of these bean samples may also contribute to the reduced ability to retain moisture (Lin et al., 2019). The OBC of raw bean flours was ∼1.4 g/g (Fig. 6B), equivalent to an OBC of 140%, which was similar to that reported in other pulse flours (Setia et al., 2019). Overall, both HT and HHP treatments did not significantly affect the OBC of bean flours. Nevertheless, there was a slight increase of OBC in HT120 samples, compared to HT15. A clear tendency could not be identified in HHP-treated samples, but in general terms, while at 5 and 15min, OBC decreased with increasing pressure, at 10 min, OBC seemed to increase slightly with pressure. Moreover, low pressure (150MPa) did not change OBC regardless of holding time, while higher pressure (300MPa and 450MPa) led to more fluctuant OBC values with increasing holding time. The variation in OBC is mainly related to the hydrophobic character of the protein (Aguilera et al., 2009). HT and HHP may affect the composition and profile of polar and non-polar aminoacids thus causing OBC fluctuations. Polar aminoacids are reported to decrease after pressure and thermal processing whereas non-polar amino acids increased in cooked beans (Kim et al., 2014; Mbithi-Mwikya, Ooghe, Van Camp, Ngundi, & Huyghebaert, 2000). A higher proportion of non-polar groups at the protein surface rather than a higher ratio of hydrophobic to hydrophilic residues alone, would be responsible for an enhanced OBC.

**Fig. 6.**
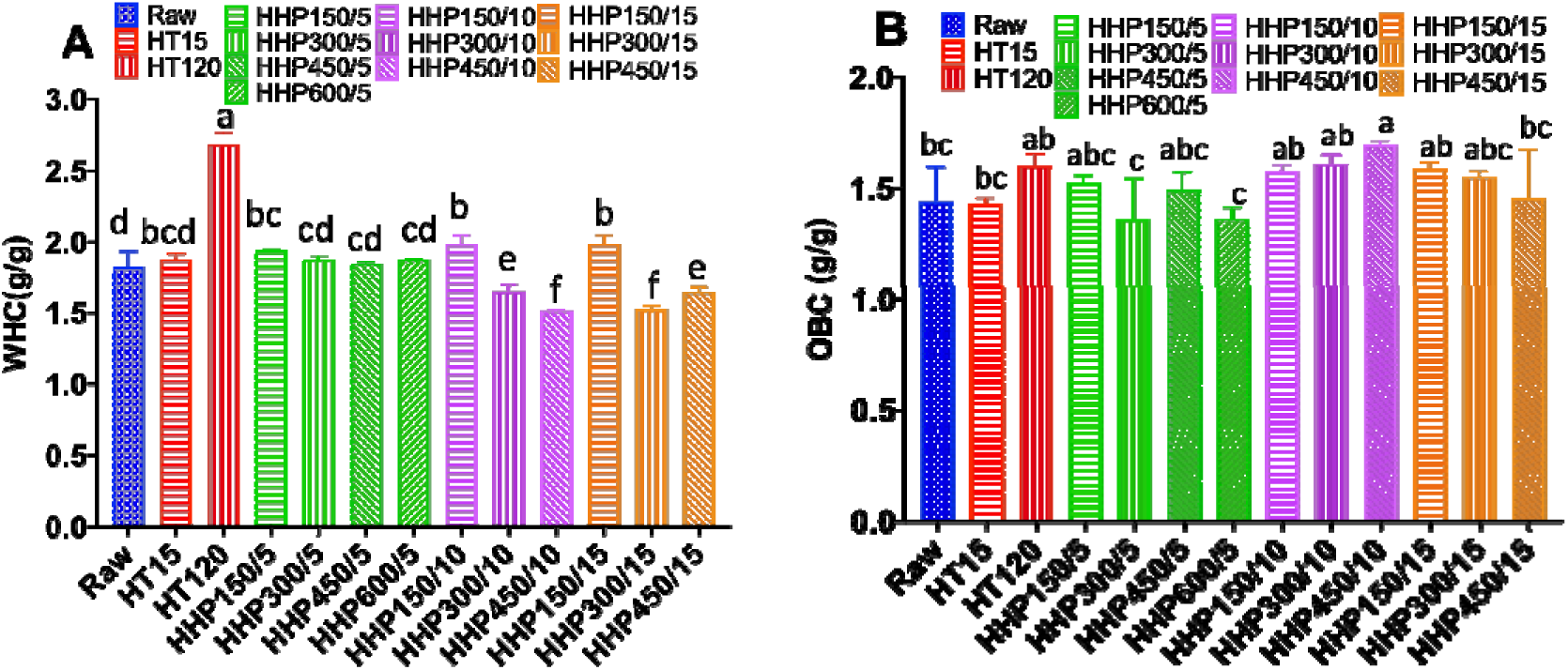
Water holding capacity (A) and oil binding capacity (B) of bean flours treated by hydrothermal and high hydrostatic pressure processing. Data are expressed as the mean values (n=3) ± standard error (SE) bar.

#### 3.4.3 Emulsifying properties

The emulsifying properties of bean flours were studied by determining the emulsifying activity index (EAI) and the emulsifying stability index (ESI) (Fig.7). The EAI indicates the ability of proteins and other surface-active molecules present in beans to adsorb to the oil-water interface and then contribute to the formation of an emulsion. The ESI indicates the stability of the adsorbed layer in a time period (Pearce & Kinsella, 1978). Raw bean flours showed good emulsifying ability with EAI ∼42 m^2^/g, higher than values reported in chickpea and lentil flours (∼20-25 m^2^/g) (Ma et al., 2011) and close to EAI of bean protein isolates (Karaca, Low, & Nickerson, 2011). HT-treatments significantly decreased the EAI of bean flours by 69-79% (Fig. 7A.) as similarly observed in thermal-treated lentils and chickpea flours. Regarding HHP, pressures<450MPa increased or maintained the EAI of bean flours when applied during 5 and 15 min. For a specific treatment time, EAI decreased with increasing pressure. As a general tendency, the decrease of EAI was proportional to the decrease of protein solubility in all processed samples (Fig. 5A), which can be expected as only the soluble protein fraction will contribute to the emulsification capacity (Kiosseoglou & Paraskevopoulou, 2011). This result is consistent with the general correlation between protein solubility and EAI previously reported in thermal-treated pulse flours (Aguilera et al., 2009). The partial protein denaturation observed at 150MPa, which result in solubilization, could have increased EAI of beans, as consequence of improved molecular flexibility and surface hydrophobicity (Damodaran & Parkin, 2017). The ESI of raw beans was about 13min (Fig. 7B), similar to the ESI of lentil and chickpea flours previously reported (Ma et al., 2011). Although ESI is attributed to the strength of the protein adsorbed layer, bean starch and fiber might also contribute to the emulsion stability by increasing the viscosity of the continuous phase, which would prevent droplet aggregation. HT and HHP treatments considerably enhanced the ESI of flours by approximately 15-17% and 13-50% respectively. As observed from these results, HT and HHP altered the protein surface activity, which is related not only to the ratio of hydrophobic/hydrophilic groups, but mostly to the protein conformation. Despite HT and HHP decreased the emulsification power of flours, both processing methods increased the ability of bean flours to stabilize the emulsion. Emulsification ability and emulsion stability seem to be affected by antagonistic molecular properties. For example, globular proteins, like globulin which represent ∼70% of bean proteins, because of more conformational constraints, adsorb slowly and only partially unfold at the interface, hence exhibiting poor emulsification power. However, since the protein retains some degree of folded structure that extends into the sub-surface promoting molecular interactions, it usually forms stable interfacial layers (Aryee, Agyei, & Udenigwe, 2018). Therefore, the decreased EAI might be due to a reduced ability of processed bean proteins to rapidly adsorb, unfold and reorient at the interface, probably because of a change in the distribution pattern of hydrophilic/hydrophobic residues on the protein surface that favored a higher proportion of hydrophilic groups at the surface. On the other hand, processing appeared to lead to protein conformational changes that enhanced interactions among neighbor proteins, resulting in the formation of a stronger, cohesive and viscoelastic layer able to better adapt to environmental changes (Aryee et al., 2018).

**Fig. 7.**
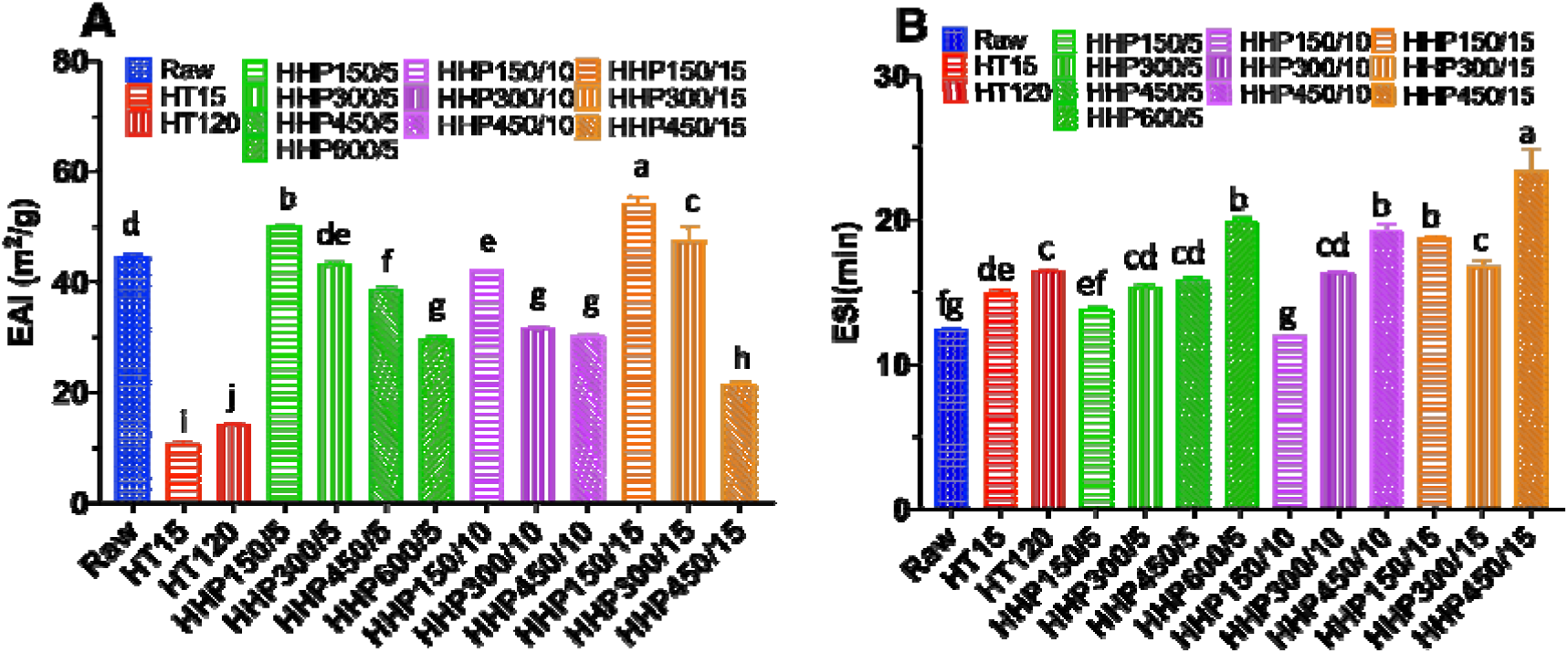
Emulsifying activity index (EAI) (A) and emulsifying stability index (ESI) (B) of bean flours treated by hydrothermal and high hydrostatic pressure processing. Data are expressed as the mean values (n=3) ± standard error (SE) bar.

## 4 Conclusions

HT and HHP processing of bean flours differently affected starch gelatinization and protein denaturation which translated into differences in gelling behavior and other functional properties. Severe HT treatment resulted in complete starch gelatinization and protein denaturation which drastically reduced the resistance to shear-thinning, protein solubility and emulsifying activity of bean flours, while increasing their water absorption capacity and cold-paste viscosity. The increased tendency to retrogradation observed in HT-treated bean flours would negatively impact the freeze-thaw stability of products to which flours are incorporated. HHP resulted in partial or no gelatinization of starch with the degree of swelling increasing with applied pressure and time. HHP-induced starch gelatinization was proportional to the increase of viscoelastic character of bean flours, and consequently, a range of gels of various strengths could be obtained. Results also showed a positive effect of HHP on pasting properties and resistance to shear-thinning and heating of flours, suggesting their suitability for batter-based products. Additionally, protein denaturation was more effectively preserved with HHP than with HT treatments resulting in superior protein solubility and emulsifying activity/stability, which may be exploited to partially replace emulsifier additives. In conclusion, compared to conventional cooking, HHP processing is a promising non-thermal technology for tailoring the functionality of bean flours and therefore for increasing their use as nutritious ingredients in a range of food formulations while reducing the time and energy required for processing and preparation.

## Acknowledgement

The authors acknowledge the funding support from the Dry Bean Health Research Program for the Northarvest Bean Growers Association. This project was also supported by the Virginia Agriculture Experiment Station and the Hatch Program of the National Institute of Food and Agriculture (NIFA), USDA. The authors thank Dr. Hengjian Wang and Mr. Brett Driver for their technical assistance with HHP treatments. We also thank ADM for providing the bean seeds.

