## Supplementary information for "Effect of thermal and high-pressure processing on the thermo-rheological and functional properties of common bean (*Phaseolus vulgaris* L.) flours"

**Table S1**

Linear regression (R^2^) of rheological models corresponding to untreated, hydrothermal (HT) and high hydrostatic pressure (HHP)-treated bean flour dispersions.

| Treatment | Herchel-Bulkley^a^ | Newtonian^b^ | Bingham^c^ | Power law^d^ |
| --- | --- | --- | --- | --- |
| **Raw** | 0.999 | 0.784 | 0.838 | 0.994 |
| **HT15** | 0.990 | 0.726 | 0.826 | 0.926 |
| **HT120** | 0.999 | 0.686 | 0.717 | 0.899 |
| **HHP150/5** | 0.998 | 0.797 | 0.922 | 0.967 |
| **HHP300/5** | 0.998 | 0.802 | 0.912 | 0.981 |
| **HHP450/5** | 0.996 | 0.705 | 0.884 | 0.987 |
| **HHP600/5** | 0.998 | 0.584 | 0.854 | 0.991 |
| **HHP150/10** | 0.998 | 0.509 | 0.877 | 0.982 |
| **HHP300/10** | 0.986 | 0.729 | 0.897 | 0.973 |
| **HHP450/10** | 0.994 | 0.754 | 0.896 | 0.974 |
| **HHP150/15** | 0.999 | 0.454 | 0.871 | 0.981 |
| **HHP300/15** | 0.997 | 0.685 | 0.872 | 0.981 |
| **HHP450/15** | 0.997 | 0.752 | 0.897 | 0.982 |

^a^Herchel-Bulkley: $\sigma=\sigma_{0}+K\dot{\gamma}^{n}$(Eq. 3)

^b^Newtontian: $\sigma=K\dot{\gamma}$ (Eq. 4)

^c^Bingham: $\sigma=\sigma_{0}+K\dot{\gamma}$ (Eq. 5)

^d^Power law: $\sigma=K\dot{\gamma}^{n}$ (Eq. 6)

$\sigma$: the shear stress (Pa); σ_0_: Yield stress (Pa); K: Consistency coefficient(Pa s^n^ ); n: Flow behavior index; $\dot{\gamma}$ : shear rate (s^-1^).

**Table S2**

| Treatment | Initial viscosity  （Pa） | Peak viscosity  （Pa） | Hot paste viscosity  （Pa） | Cold paste viscosity  （Pa） |
| --- | --- | --- | --- | --- |
| **Raw** | 530±15^d^ | 9970±299^c^ | 1869±56^h^ | 9736±292^d^ |
| **HT15** | 33325±999^b^ | - | 15243±457^a^ | 59122±1172^a^ |
| **HT120** | 39192±1175^a^ | - | 7954±238^c^ | 22375±627^b^ |
| **HHP150/5** | 63±2^d^ | 7599±228^ef^ | 4803±144^e^ | 2301±62^f^ |
| **HHP300/5** | 21±0.6^d^ | 7629±220^ef^ | 3374±101^g^ | 156±5^g^ |
| **HHP450/5** | 23±0.7^d^ | 8576±215^d^ | 6196±185^d^ | 32±1^g^ |
| **HHP600/5** | 1844±55^c^ | 19222±527^a^ | 11962±358^b^ | 15101±343^c^ |
| **HHP150/10** | 208±6^d^ | 11476±344^b^ | 4649±139^e^ | 5591±126^e^ |
| **HHP300/10** | 63±2^d^ | 6993±209^f^ | 4507±135^e^ | 30±1^g^ |
| **HHP450/10** | 22±0.7^d^ | 7971±213^de^ | 6282±127^d^ | 122±4^g^ |
| **HHP150/15** | 45±1^d^ | 8052±241^de^ | 3829±188^fg^ | 7053±121^e^ |
| **HHP300/15** | 36±2^d^ | 7909±237^de^ | 4190±114^ef^ | 60±2^g^ |
| **HHP450/15** | 18±0.5^d^ | 7370±221^ef^ | 5917±125^d^ | 37±1^g^ |

Effect of hydrothermal and hydrostatic pressure treatments on the pasting parameters of bean flours
